# Computational Simulations of the 4-D Micro-Circulatory Network in Zebrafish Tail Amputation and Regeneration

**DOI:** 10.1101/2021.02.10.430654

**Authors:** Mehrdad Roustaei, Kyung In Baek, Zhaoqiang Wang, Susana Cavallero, Sandro Satta, Angela Lai, Ryan O’Donnell, Vijay Vedula, Yichen Ding, Alison Lesley Marsden, Tzung Hsiai

## Abstract

Wall shear stress (WSS) in the micro-vasculature contributes to biomechanical cues that regulate mechanotransduction underlying vascular development, regeneration, and homeostasis. We hereby elucidate the interplay between hemodynamic shear forces and luminal remodeling in response to vascular injury and regeneration in the zebrafish model of tail amputation. Using the transgenic *Tg*(*fli1:eGFP; Gata1:ds-red*) line, we were able to track the enhanced green-fluorescent protein (eGFP)-labeled endothelial lining of the 3-D microvasculature for post-image segmentation and reconstruction of fluid domain for computational fluid dynamics (CFD) simulation. At 1 day post amputation (dpa), dorsal aorta (DA) and posterior cardinal vein (PCV) were severed, and vasoconstriction developed in the dorsal longitudinal anastomotic vessel (DLAV) with a concomitant increase in WSS in the segmental vessels (SV) proximal to the amputation site and a decrease in WSS in SVs distal to amputation. Simultaneously, we observed angiogenesis commencing at the tips of the amputated DLAV and PCV where WSS was minimal in the absence of blood flow. At 2 dpa, vasodilation occurred in a pair of SVs proximal to amputation, resulting in increased flow rate and WSS, whereas in the SVs distal to amputation, WSS normalized to the baseline. At 3 dpa, the flow rate in the arterial SV proximal to amputation continued to rise and merged with DLAV that formed a new loop with PCV. Thus, our CFD modeling uncovered a well-coordinated micro-vascular adaptation process following tail amputation, accompanied by the rise and fall of WSS and dynamic changes in flow rate during vascular regeneration.

## Introduction

Biomechanical forces modulate vascular morphological adaptation during development (1, 2). While mechanotransduction mechanisms underlying arterial inflammatory responses and oxidative stress are well-elucidated in the macro-scale in association with high Reynolds number flow (3), much less is known about the interplay between hemodynamic forces and vascular remodeling in the micro-scale in association with low Reynolds number flow (4, 5). While endothelial dysfunction in the microvasculature is well-recognized in myocardial ischemia, diabetes, and peripheral artery diseases (PAD) (6–8), investigation into vascular injury-mediated changes in hemodynamic forces and adaptive vascular remodeling in the microvasculature remained an experimental challenge.

Zebrafish (*Danio Rerio*) embryos are transparent for light microscope imaging and genetically tractable for studying shear stress-mediated myocardial and vascular development (9). We previously established 4-D light-sheet fluorescent microscopy (LSFM) to study the development of cardiac trabeculation during zebrafish cardiac morphogenesis (10, 11). Combining LSFM and computational fluid dynamics (CFD), we have provided biomechanical insights into wall shear stress (WSS)-mediated endocardial trabeculation and outflow tract valvulogenesis in embryonic zebrafish (12–14). Thus, we sought to use the transgenic zebrafish *Tg(fli1:eGFP; Gata1:ds-red)* line, in which enhanced green fluorescent protein-labeled endothelium allows for segmentation of 3-D vasculature and *ds-red*-labeled blood cells (mainly erythrocytes) for CFD reconstruction and micro particle image velocimetry (PIV).

Wall shear stress (WSS) intimately regulates mechanotransduction underlying vascular development, repair, and homeostasis (15, 16). The spatial and temporal variations in fluid shear stress in the segmental vessels (SVs) of zebrafish embryos modulate the transformation of arterial to venous vessels, leading to a balanced distribution of arterial and venous SVs during vascular development (17). In this context, the zebrafish microvascular network can be considered as an circuit analogy model consisting of the dorsal aorta (DA), segmental vessels (SVs), and the posterior cardinal vein (PCV). This model allows for simulating the microvascular response to blood cell partitioning in each SV, leading to uniform flow distribution (18, 19). However, this zero-dimensional resistance modeling neglects bidirectional flow in the dorsal longitudinal anastomotic vessel (DLAV) network, vascular remodeling, including constriction and dilation, and angiogenesis of the DLAV and posterior cardinal vein (PCV) to form a new loop following tail amputation. Zero-dimensional modeling approaches also overlook the local variations in vessel diameter in association with the hemodynamic resistance in the microvasculature. For this reason, our CFD modeling sought to account for hemodynamic resistance for accurate estimation of WSS and luminal interaction.

We hereby demonstrate a 3-D CFD model encompassing 4 time-lapse stages of vessel regeneration following tail amputation, and subsequent loop formation between DA and PCV. We employed *eGFP*-labeled vascular endothelium by sub-micron confocal imaging and *ds-Red*-labeled blood cells to reconstruct the computational model and to predict the interplay between hemodynamic changes vascular morphological remodeling. We discovered a coordinated vascular adaptation to tail amputation that disconnected the DA from the PCV, followed by DLAV vasoconstriction at 1 day post amputation (dpa), and SV dilation at 2 dpa, coupled with the rise and fall of WSS and the hemodynamic changes in flow rates. At 3 dpa, flow rate in the arterial SV proximal to amputation continued to rise and merged with DLAV to form a new loop with PCV. Thus, elucidating the interplay between shear forces and luminal remodeling provides new biomechanical insights into the micro-scale hemodynamics, with translational implications in vascular injury and repair under low Reynolds number flow in diabetes and PAD.

## Materials and Methods

Zebrafish (*Danio Rerio*) has the capacity to regenerate following distal tail amputation, providing a genetically tractable model to study changes in microcirculation in response to vascular development, injury, and regeneration (20–22). We performed tail amputation on embryonic zebrafish to assess the dynamic changes in wall shear stress (WSS) and in flow rates in response to microvascular adaptation and remodeling proximal to the amputation site. The investigation of hemodynamic changes and morphological remodeling was established by imaging the transgenic *Tg(fli1:eGFP; Gata1:ds-red)* zebrafish embryos from 4 to 7 days post fertilization (dpf) (**Fig. 1a–c**), followed by post-imaging segmentation (**Fig. 1d, e**), and 3-D computational fluid dynamics (CFD) simulation (**Fig. 1g, h**). The enhanced green fluorescent protein (*eGFP*)-labeled endothelial cells expressed vascular endothelial growth factor (VEGF) receptor, denoted as *fli1*, and the *ds-red*-labeled blood cells (mostly erythrocytes and other blood lineages) allowed for tracking their movement. Thus, sequential analyses from stitching of the 3-D confocal imaging to computational modeling to particle imaging velocity (PIV) validated the hemodynamic changes proximal to the amputation site, vascular constriction and dilation, and new vessel formation (angiogenesis) in response to tail amputation (**Fig. 1**).

**Fig. 1.**
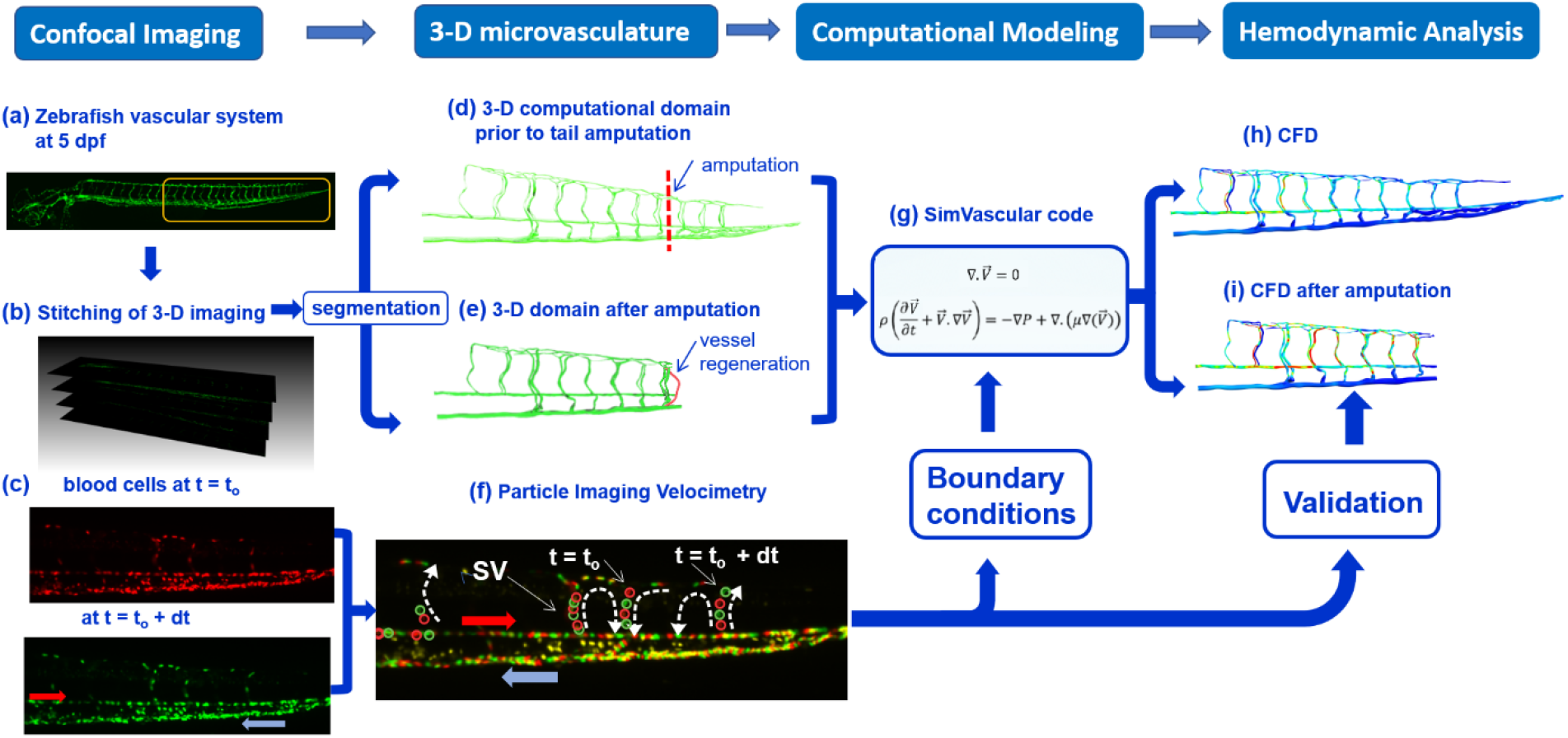
A pipeline of post-imaging segmentation, simulation, and validation for computational fluid dynamics (CFD). (a) Confocal microscopy was used to image the vascular system from a transgenic *Tg* (*fli:eGFP*) zebrafish embryo at 5 days post fertilization (dpf). The vasculature was visualized via the enhanced green fluorescent protein (*eGFP*)-labeled endothelial cells that express VEGF receptor, *fli1*. (b) Stitching of the 3-D confocal images provided reconstruction of the vasculature for post-imaging segmentation. (c) The *ds-red* labeled blood cells (*gata1:ds-red*) in the zebrafish tail microvasculature were monitored at two different time points. The motion of *ds-red*-labeled blood cells allowed for quantitative assessment of the blood flow at t = t_o_ and t = t_o_ + dt in the microvascular network. Following segmentation, 3-D computational domain was reconstructed (d) prior to and (e) after tail amputation. (f) Particle image velocimetry (PIV) was performed to track the individually *ds-red*-labeled blood cells in the segmental vessels (SV) from t = t_o_ to t = t_o_ + dt. (g) SimVascular open-source blood flow modeling software was employed to solve the Navier-Stokes equations for incompressible flows, and boundary conditions were derived from PIV. (h) CFD analyses revealed wall shear stress prior to and after tail amputation. The magnitude of velocity in SVs was validated by PIV.

### Transgenic Zebrafish Model

The transgenic *Tg(fli1:eGFP; Gata1:ds-red)* embryos were rinsed with fresh standard E3 medium and cultivated at 28.5°C for 4 days after fertilization. Standard E3 medium was supplemented with 0.05% methylene blue (Sigma Aldrich, MO) to prevent fungal outbreak and 0.003% phenylthiourea (PTU, Sigma Aldrich, MO) to reduce melanogenesis. At 4 days post fertilization (dpf), embryos were randomly selected for immobilization with neutralized tricaine (Sigma Aldrich, MO) to undergo tail amputation. Zebrafish responses to anesthesia were monitored by the movement of the caudal fin. The tail was amputated at ~100 *μm* from the distal end. Following amputation, embryos were returned to fresh E3 medium and maintained for 3 days.

### Post-amputation Imaging of Blood Flow and Microvasculature

The images of blood cells (*Gata1:ds-Red*) transport were captured by the inverted microscope (Olympus, IX70) and digital CCD camera (QIclick, Teledyne Qimaging, Canada). We performed time-lapsed imaging from 4 dpf to 7 dpf or 0 to 3 7dpa for both the amputated and control fish (amputation = 7; control = 5). The acquired images revealed the motion of erythrocytes in the microvasculature throughout multiple cardiac cycles.

The transgenic zebrafish *Tg(fli1:eGFP; Gata1:ds-red)* line was employed to capture *GFP*-labeled endothelium lining of the microvasculature undergoing remodeling following amputation from 4 to 7 dpf. The confocal-imaging system (Leica TCS-SP8-SMD) provided the isotropic submicron resolution in 3 dimensions (~500 nm in-plane and in *z* direction) for segmentation of the 3-D microvasculature. To reach this resolution, we stitched the high magnification (64*x* objective lens) images at 4-7 dpf or 0-3 dpa, and we used 5 dpf for segmentation of the tail microvascular network.

We used our in-house light-sheet for the time-lapse imaging of zebrafish tail to assess the vascular remodeling in response to amputation. Our system utilized a continuous-wave laser (Laserglow Technologies, Canada) for planar illumination and a detection module including a set of filters and scientific CMOS (sCMOS, ORCA-Flash4.0, Hamamatsu, Japan). Our system provides a resolution of 2*μm* that was adequate for the measurement of microvascular diameter variation after amputation. (23)

### Segmentation of Microvascular Network

We used the 3-D *GFP*-labeled images of endothelial lining to reconstruct the computational domain in SimVascular (**Fig. 1b**) (24). The pathline of each vessel was produced by connecting the points in the center of the lumen along the vessel in the 3-D image stack. For each vessel, the cross-section was calculated by interpolation of the manually specified points on the wall/lumen interface (1*D* → 2*D*). Linking the cross-section areas along the vessel path with a spline function, we reconstructed each vessel in the network (2*D* → 3*D*). Overall, we generated the vascular network for 21 segmental vessels (SVs), 20 dorsal longitudinal anastomotic vessels (DLAVs), dorsal aorta (DA), posterior cardinal vein (PCV), and capillary venous plexus (**Fig. 1d, e**).

We implemented the post-amputation remodeling in microvasculature on the 3-D reconstructed computational domain (Fig. 3 a–c). At 1 dpa, we removed 8 SVs (6 SVs for amputation and 2 SVs for vasoconstriction) from the zebrafish tail microvascular network distal to the heart to model the microvascular response to amputation followed by vasoconstriction at the DLAV (1 dpa). At 2 dpa, 2 SVs were added back to the microcirculation network where vasodilation was implemented (see Fig. 3 d, e). We employed the time-lapse light-sheet fluorescent microscopy images to measure the average length and diameter of the regenerated vessel. The arterial SVs proximal to amputation was connected to the PCV to model loop formation at 3 dpa (Fig. 3 c).

### Particle Image Velocimetry for the Boundary Conditions and Validation

We developed and implemented our PIV code on *ds-Red*-labeled blood cells. To determine the boundary conditions for the inlet (DA) and to validate the results, we measured the velocity in DA and individual SVs. Our custom-written PIV algorithm was able to correlate with the intensity distribution of *ds-Red-*labeled erythrocytes in different time frames. A 2-D continuous polynomial function was fitted to the correlation distribution, and the position of the maximum correlation was determined from the polynomial function between two timeframes. At 20 frames per second, the computed erythrocyte position allowed for estimating the velocity in DA and SVs (**Fig. 1c, f**).

### 3-D Computational Modeling

A stabilized 3-D finite element method (FEM) was applied for CFD analysis to simulate the blood flow in the microvascular network using SimVascular’s svFSI solver (24). The Navier-Stokes equations were solved in svFSI with an incompressible flow assumption to quantify hemodynamic changes in response to tail amputation and vascular regeneration from 0 to 3 dpa (4-7 dpf) (**Fig. 1g**). In the setting of low Reynolds number (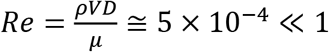, where D is defined as the vessel diameter) in the microvascular network, viscous forces dominate over inertial forces, and the diameters and lengths of the vessels governed the pressure distribution and flow rates. Due to the low Reynolds number (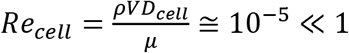, where D is defined as the erythrocyte diameter), the pathway of the *ds-Red*-labeled erythrocytes followed the streamline of blood flow. Unlike the anucleate red blood cells in the mammalian circulation, zebrafish erythrocytes contain nuclei, and are thus, less deformable.

Despite the time-dependent nature of blood flow in zebrafish microcirculation, the Womersley number (ratio of transient inertial force to viscous force: 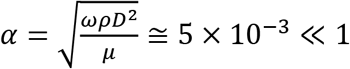) was small, and the time-dependent terms in the Navier-Stokes equations were neglected in our continuum modeling (24).

We calculated the inlet velocity into DA during a cardiac cycle using the PIV code, and the time-averaged magnitude of *V*_ave_ = 238 μm·s^−1^ was applied as the inlet boundary conditions. Given the small Womersley number (*α* ≪ 1), a fully developed parabolic flow was considered in the inlet of DA (red arrow in **Fig. 1f**). A zero-pressure boundary condition was applied at the outlet of PCV (blue arrow in **Fig. 1f**). A non-slip boundary condition was assumed on the vessel walls.

To incorporate non-Newtonian effects of flow in the micro-circulation, we used a non-Newtonian viscosity model of the form:

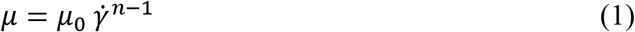

where *μ* is blood viscosity, 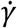 the shear rate, *μ*_0_ the consistency index, and *n* the power-law index. We employed μ_0_ = 1.05 × 10^−3^ *Pa*.*s^n^* and *n* = 1.09 for the power-law equation (25). We fitted the parameters and implemented this model into a Carreau-Yasuda model in svFSI (26). We further analyzed the non-Newtonian effects on the flow rates and wall shear stress (WSS) in SVs by performing a parametric study based on the power-law index (*n*).

## Results

### Segmentation of 3-D microvasculature

Following 3-D stitching of the confocal images from the intact and amputated zebrafish tail, we performed segmentation of the microvascular network (**Fig. 2.a–b**). Next, 3-D computational domain was reconstructed to simulate hemodynamic changes in response to tail amputation and vascular regeneration. The square box indicates the region of interest (ROI) for vascular segmentation and simulation (**Fig. 2.a**). A representative 3-D computational domain was acquired from the segmentation process (**Fig. 2.c–d**). The pressure drop across the venous plexus was deemed to be negligible due to both the short vessel length and low hemodynamic resistance. These considerations allowed for connecting the venous SV with the PCV for segmentation through the largest venous plexus.

**Fig. 2.**
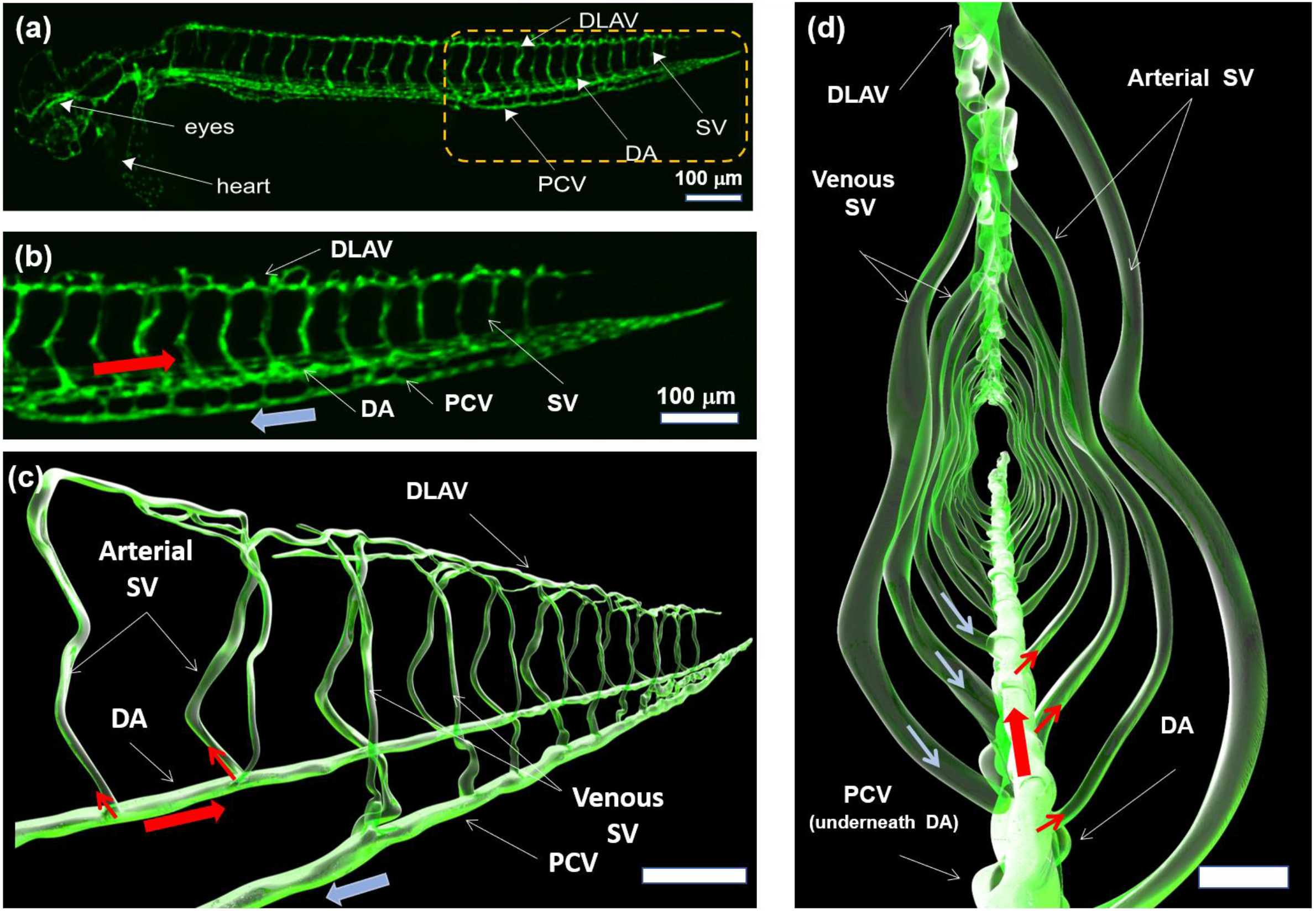
(a) Transgenic Tg(*fli1:GFP*) zebrafish line was used to visualize the 3-D vascular network. The dashed rectangle (yellow) indicates the tail region from which vascular network was reconstructed for CFD simulation. (b) Magnification of the tail region depicts the flow direction in the DA in relation to PCV. (c) Representative segmentation of the 3-D vasculature recapitulates DA, SV, DLAV, and PCV. (d) Magnification of the yellow rectangle reveals the cross-section of the trail region where the vascular connectivity occurs between DLAV and PCV and between DA and SVs. Note that arterial SVs branch off from the DA, whereas venous SVs drain into PCV. CFD: computational fluid dynamics, DA: dorsal aorta, SV: segmental vessels, DLAV: dorsal longitudinal anastomotic vessel, and PCV: posterior cardinal vein.

While DLAVs are commonly assumed to be a single straight vessel for unidirectional flow, bidirectional flow occurs in the DLAVs in parallel (**Supplementary Video S1-S2**). For this reason, a parallel pattern was proposed for the DLAVs according to their interconnection in the reconstructed vascular network (**Fig. 3.b and Supplementary Fig. S1**). Thus, we demonstrate a pair of DLAVs in parallel, intersecting with the arterial and venous SV (in blue and red) to regulate the blood flow between SVs.

**Fig. 3.**
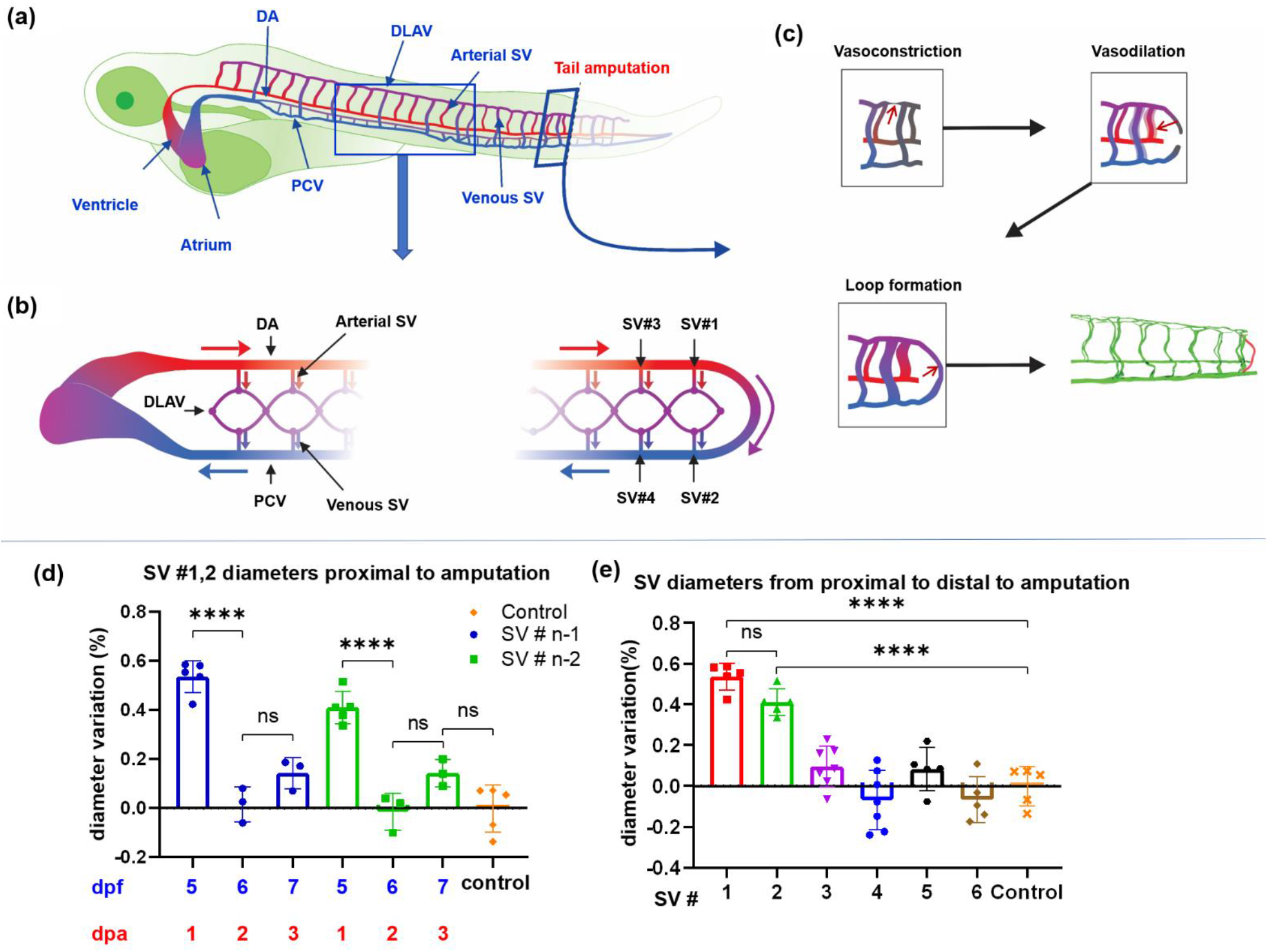
Zebrafish embryonic cardiovascular circulatory system in response to tail amputation and regeneration. **(a)** Arterial circulation is denoted in red and venous in blue. Two regions were magnified to reveal vascular network; namely, top view and tail amputation. (**b**) The top view reveals that DA originates from the ventricle, and PCV returns to the atrium. Arterial SV connect with DLAV, which drain to PCV via venous SV. **(c)** The tail amputation site (side view) illustrates SVs undergoing vasoconstriction, vasodilation, and loop formation proximal to amputation at 1, 2, 3 dpa. The proximal SVs were numbered from #1 to #6. **(d)** The changes in proximal SV diameters were compared from 0 dpa to 3 dpa. Both SV#1 and SV#2 underwent significant changes in diameter due to vasoconstriction at 1 dpa and vasodilation at 2 dpa (p < 0.05 vs. control, n=5). The changes at 3 dpa were statistically insignificant. **(e)** Changes in diameters were significant to the proximal SV#1 and SV#2 as compared to the distal SV#3-6. ns: not significant, DA: dorsal aorta, SV: segmental vessels, DLAV: dorsal longitudinal anastomotic vessel, and PCV: posterior cardinal vein, dpa: days post amputation.

### Vascular Remodeling Proximal to Tail Amputation Site

Confocal images revealed remodeling of the vessels and changes in blood flow from 0 to 3 days post-amputation (**Fig. 3**). At 0-1 dpa, DLAVs underwent vasoconstriction, accompanied by a reduction in blood flow in the SVs proximal to the amputation site while both DLAV and PCV started to form a new connection toward the tail region (**Supplementary Fig. S2 and Video S3-S4**) (27). At 0-1 dpa, SVs proximal to the amputation site (SV # a+1, a+2, where *a* denotes the number of SVs that were removed after amputation) underwent statistically significant vasodilation (SV # a+1 and SV # a+2, *p* < 0.0001, n=61), whereas the diameter variations in the SVs distal from the amputation site (SV # > a+3) remained statistically unchanged (SV # a+3 to # a+6, *p* > 0.05, n= 61) (**Fig. 3.d–e, Supplementary Video S4**). At 2 dpa, the DLAV proximal to SV# a+1 started to form a new loop with PCV (**Fig. 3.c and Supplementary Video S5-S6**). At 3 dpa, vasodilation in SV # a+1 and # a+2 attenuated in conjunction with the complete loop formation between DLAV and PCV, allowing for vascular regeneration to restore blood flow in the tail (**Fig. 3.c and Supplementary Video S7-S8**). Thus, we unveil micro-circular remodeling proximal to the tail amputation site, where the diameters of DLAV reduced at 1 dpa, followed by increased diameters in arterial SV # a+1 and # a+2, with concomitant angiogenesis in DLAV and PCV to form a new loop at 2 dpa, followed by normalization of SV diameter when a new loop connected the arterial and venous circulation.

### Particle Image Velocimetry to Validate CFD

To validate our CFD model for low Reynolds number flow in the tail region, we performed particle image velocimetry (PIV) to obtain the mean velocity of blood flow in the SVs from proximal to distal to the amputation site. CFD simulation revealed velocity ranging from 130 to 550 μm·s^−1^, which was comparable to that obtained by PIV (**Fig. 4.e**). The CFD results deviated from those of PIV by ~8%, and the velocity magnitude of the most distal SV #15 was significantly higher than that of the proximal SVs, indicating a good agreement with the experimental data.

**Fig. 4.**
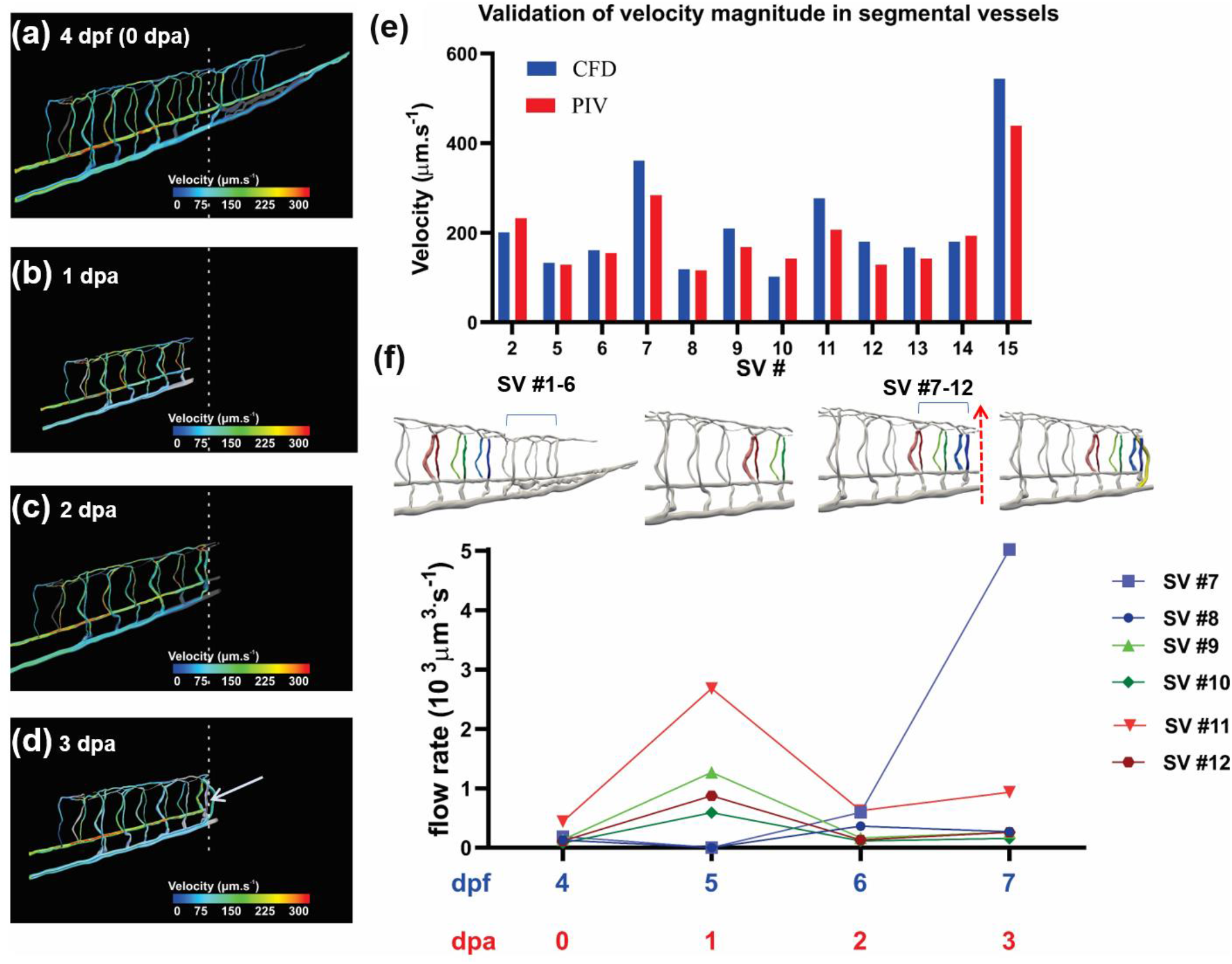
Computational Fluid Dynamics (CFD) simulates changes in hemodynamic responses to tail injury and regeneration. Velocity streamlines in the computational domain were color-coded by the velocity magnitude (a) at 4 days post fertilization (dpf) prior to tail amputation or 0 days post amputation (dpa), (b) 1 dpa, (c) 2 dpa, and (d) 3 dpa. Arrow indicates vascular repair with a loop formation. (e) Variations in velocity from the proximal to distal segmental vessels (SV) were compared between CFD and particle imaging velocimetry (PIV) prior to tail amputation. (f) The dynamic changes in flow rates developed from the proximal to distal SVs before (4 dpf or 0 dpa) and after tail amputation (5-7 dpf or 1-3 dpa). Note that flow rate in SV #7 increased at 3 dpa when DLAV formed a loop with PCV.

### Changes in the Hemodynamic Parameters Prior to Tail Amputation

Results from the 3-D CFD simulations included wall shear stress (WSS), pressure gradients, velocity profiles, and flow rates in the individual SVs. Our model verified the unique pattern of blood flow that recirculates between two SVs in parallel (**Supplementary Video S1-S2**). The blood velocity and flow rate in DA decreased toward the tail region where arterial blood flow drains into the SVs and DLAV. Thus, this compartmental increase in hemodynamic resistance in both the SVs and DLAVs toward the tail region results in a decrease in flow rate in the DA.

Our 3-D simulations further demonstrated the bidirectional flow in the DLAVs in parallel (see **Fig. 3b and Supplementary Video S2**). The flow direction in each DLAV was regulated by the pressure difference between the arterial and venous SVs corresponding to the diameter distribution in SVs under Stokes flow (*Re* ≅ 10^−3^). As a result, the combination of DLAVs in parallel and SVs of various diameters modulated the flow rates and WSS (**Fig. 5.d**). The flow rate in each SV was primarily regulated by the changes in diameters of the arterial and venous SVs and the interconnecting DLAV (**Fig. 4.d, f**). Taken together, our *in-silico* model predicted that small changes in the arterial SV diameter and its branching vessels regulate the direction of blood flow in the micro-circular network.

**Fig. 5.**
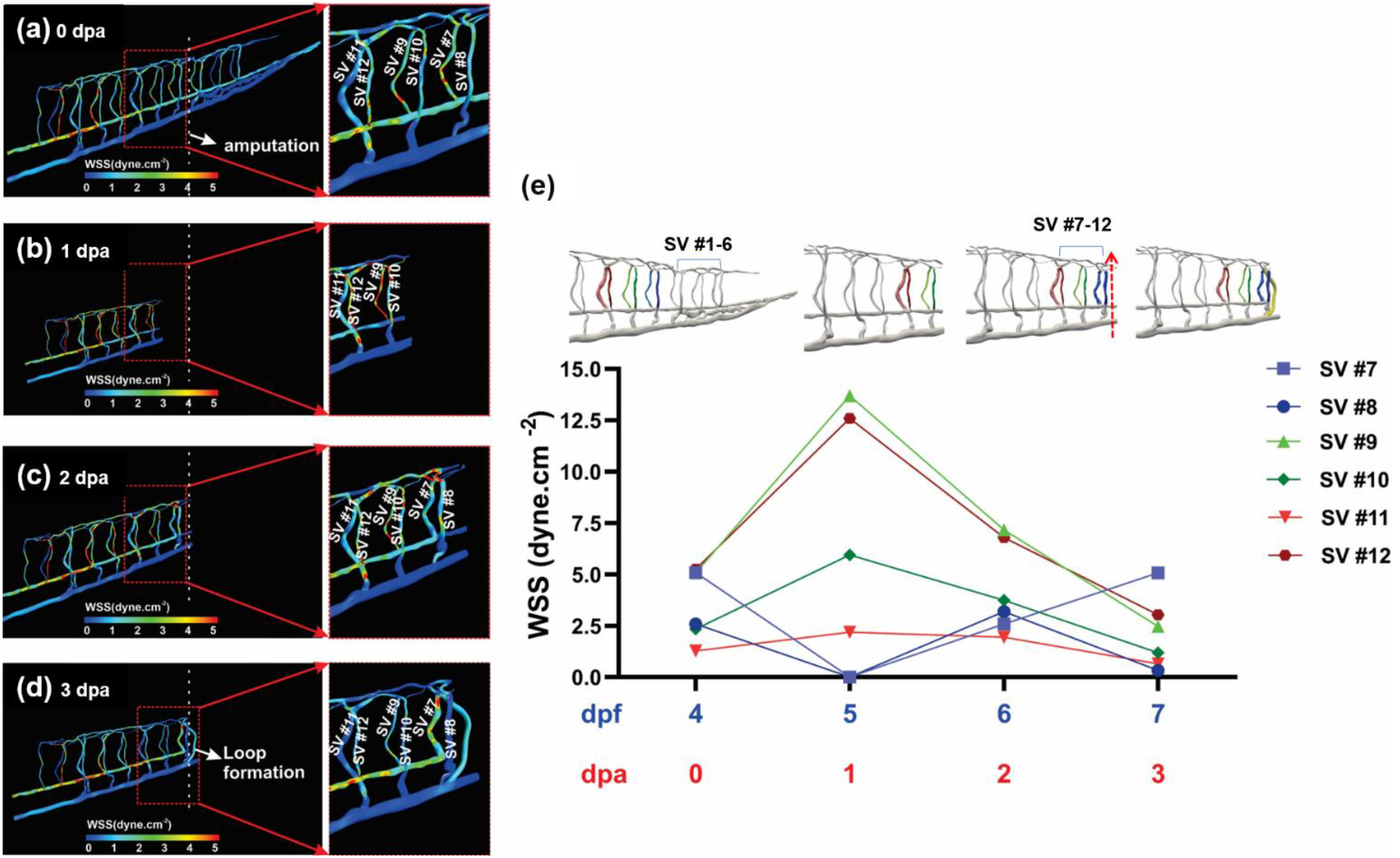
Computational fluid dynamics (CFD) simulated changes in wall shear stress (WSS) in response to tail amputation and regeneration. (a) Relatively high WSS developed in the distal arterial segmental vessels (SV) at 4 days post fertilization (dpf) prior to tail amputation. (b) An increase in WSS occurred in the proximal SV following tail amputation at 1-2 days post amputation (dpa). (c) An increase in WSS developed in the dorsal longitudinal anastomotic vessel (DLAV) and arterial SV proximal to amputation site, along with a reduction in WSS in the rest of SVs at 2 dpa. (d) A reduction in WSS developed in distal SVs in response to a loop formation connecting DLAV with posterior cardinal vein (PCV) at 3 dpa. (e) The mean WSS from the distal SV were compared prior to and post tail amputation, followed by loop formation at 3 dpa.

### Changes in WSS in Response to Tail Amputation and Regeneration

We compared changes in flow rates and WSS in the vascular network from 4 dpf (pre-amputation) to 7 dpf (3 dpa - loop formation) (**Fig. 4**). Next, we compared the changes in the area-averaged WSS (mean WSS) and flow rates in SVs following tail amputation (**Fig. 5.b–c**). At 0 to 1 dpa, we used the transgenic Tg (*fli1:eGFP*) zebrafish line where vascular endothelial growth factor (VEGF) receptors expressed in the vascular endothelial cells were encoded with the enhanced green fluorescent protein. We were able to visualize the DLAVs proximal to the amputation site undergoing vasoconstriction, restricting blood flow, along with a reduction in WSS in SVs #7-8 (**Supplementary Fig. S2 and Video S2**). Our simulation further showed that WSS increased following tail amputation in the rest of SVs between 0 to 1 dpa (**Fig. 5.a–e**). This increase was preferentially higher in the arterial SVs (i.e. SV #9, 12) than in the venous SVs (i.e. SV #10, 11) as the former experienced the pressure variation upstream from the DLAV (**Fig. 5.e**). At 0 - 1 dpa, the eGFP-labeled images revealed that DLAV and PCV formed 75% of the new vessel length while proximal SVs (SV #7, 8) developed a relatively low WSS due to DLAV vasoconstriction and the average WSS in the rest of the SVs was elevated comparing to unamputated fish (**Fig. 5.a-e**).

Simulation of blood flow at 2 dpa predicted that vasodilation of the proximal SVs (SV #7-8), and subsequent new loop connecting DLAV with PCV resulted in a proximal increase in flow rates and normalization in WSS in vascular network to the range prior to amputation (**Fig. 5. c, e and Supplementary Video S5**). However, WSS and flow rates in the distal SVs decreased at 2 dpa to forestall further SV dilation (**Fig. 5.c, e**). Our simulation also predicted that vasodilation increased the flow rate (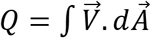, where *Q* is flow rate, V is velocity and A is cross sectional area) proximal to the amputation site (SV #7-8) while the reduction in WSS further attenuated SV dilation (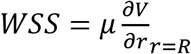, where *μ* is viscosity, and *r* is the radius). Thus, our model demonstrates that SV dilation mediated a reduction in WSS (**Fig. 3.f and Fig. 5.e**).

The CFD model captured the loop formation of the regenerated vessel occurring distal to the arterial SV proximal to the amputation site (SV# 7). This SV supplied the arterial flow to DLAV, which underwent angiogenesis to form a new loop with PCV (**Fig. 4.d, Fig. 5.d**). This flow path (DA→SV #7→DLAV→regenerated vessel→PCV) decreased the blood supply to the rest of the SVs and DLAVs, leading to the DA→PCV loop formation, and an increase in flow rates and WSS in the arterial SV proximal to amputation (**Fig. 4.d, f and Fig. 5.d, e**). However, the blood flow to the venous SV proximal to the amputation site was reduced by the new loop between DLAV and PCV, resulting in a reduced WSS (**Fig. 4.d, f and Fig. 5.d, e**). Overall, CFD simulation demonstrated the changes in flow rate and WSS in response to the variation in SV diameters and flow rates needed to restore micro-circulation following tail amputation and regeneration.

### The Non-Newtonian Effects on the Flow Rates and Wall Shear Stress

From the power law model, we assessed the effects of non-Newtonian index on the microcirculation. We normalized the results according to the reported value for non-Newtonian index (*n* = 1.09). The normalized mean WSS in SVs were compared with different non-Newtonian indexes (**Fig. 6.a, c**). Our parametric study indicates a non-linear effect of non-Newtonian index on WSS, with *n* = 1.09 indicating a maximum WSS condition (**Fig. 6.c**). Similarly, the flow rates in SV exhibited a comparable trend (**Fig. 6.b**), and the overall circulation of blood was independent of the non-Newtonian behavior.

**Fig. 5.**
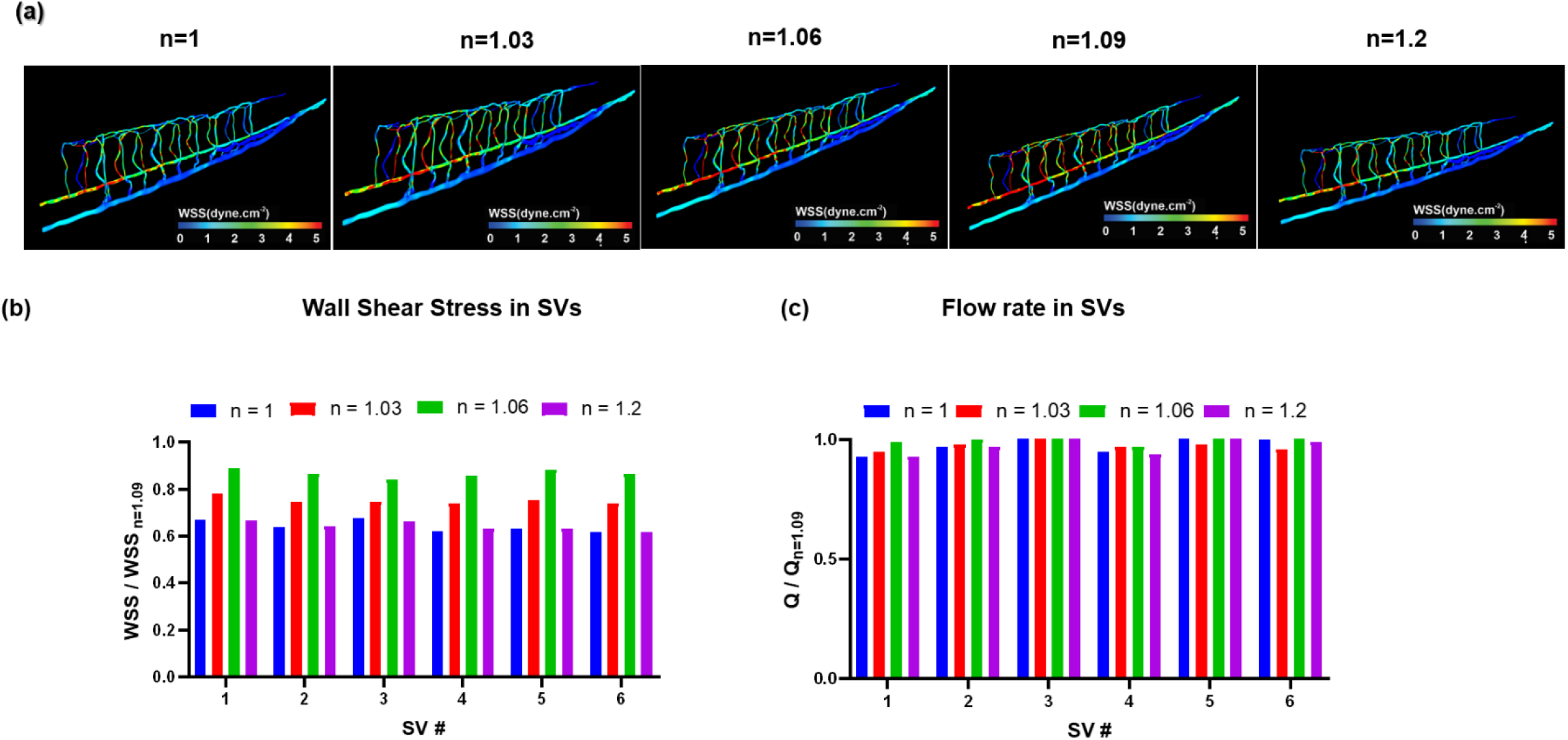
The effects of non-Newtonian Index (n) from 1 to n =1.2. (a) The non-dimensionalization was performed using the experimental index at n =1.09. Variations in WSS and flow rates were non-linear. The highest value for WSS was obtained using standard value n=1.09. (a) Dimensionless wall shear stress and (b) flow rates were compared from proximal to distal SVs.

We compared WSS contours for values of *n* from 1 to 1.2 (**Fig. 6.a**). The overall blood flow to the microvasculature was not altered by varying the non-Newtonian model indices.

## Discussion

Fluid shear stress imparts both metabolic and mechanical effects on vascular endothelial function (3). During development, hemodynamic forces such as shear stress are intimately linked with cardiac morphogenesis (28). We have demonstrated that peristaltic contraction of the embryonic heart tube produces time-varying shear stress (∂τ/∂t) and pressure gradients (∇*P*) across the atrioventricular canal in a transgenic zebrafish *Tg*(*fli-1:eGFP; Gata1:ds-red*) model of cardiovascular development (1). The advent of the zebrafish genetic system has enabled the application of *fli1* promoter to drive expression of enhanced green fluorescent protein (eGFP) in all vasculature throughout embryogenesis (29); thereby, allowing for 3-D visualization for CFD simulation. By integrating 3-D reconstruction of confocal images with CFD simulation, we unraveled the interplay between wall shear stress (WSS) and microvascular adaptation in response to tail amputation and vascular regeneration. Tail amputation resulted in disconnection between DA and PCV, that triggered a well-coordinated DLAV vasoconstriction, accompanied with the initial decrease in WSS and flow rate in the SVs proximal to the amputation site. Simultaneously, an increase in WSS and flow rate in the SVs distal to amputation site developed due to bypass of the blood from DA during 0-1 dpa. WSS normalized to baseline in the SVs during vasodilation at 2dpa when DLAV and PCV formed a new loop at 3 dpa, whereas flow rates in arterial SV merged with DLAV and continued to rise. Thus, our data provide the new hemodynamic insights in micro-circular network injury and repair at low Reynolds numbers.

Hemodynamic shear forces are well-recognized to modulate endothelial homeostasis (30, 31), migration (32), vascular development and regeneration (33). At a steady state, shear stress (*τ*) is characterized as dynamic viscosity (*μ*) of a fluid multiplied by its shear rate 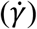, defined as the tangential velocity gradient (3): 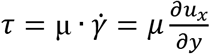. The zebrafish system allows for genetic manipulations of viscosity to alter shear stress (12, 22, 34). We previously reported that injection of *Gata*-1a oligonucleotide morpholino (MO) to inhibit erythropoiesis resulted in a decrease in viscosity-dependent shear stress, and subsequent impaired vascular regeneration following tail amputation (35). After *TNNT-2* MO injection to inhibit myocardial contraction and blood flow, zebrafish embryos also failed to develop tail regeneration at 3 dpa, whereas injection of *erythropoietin (Epo)* mRNA activates erythrocytosis to increase viscosity, resulting in restored vascular regeneration (35). Thus, the rise and fall of WSS in adaptation to vascular remodeling in SV and DLAV proximal to the amputation site occurred during the formation of a new loop connecting between DLAV and PCV.

Our computational modeling was developed from the dynamic adaptation of vessel morphology to tail injury and vascular regeneration in the micro-scale under low Reynolds numbers. Choi *et al*. studied the variation in WSS in an embryonic zebrafish tail using an circuit analogy model (19), and they demonstrated changes in WSS in the microvascular network by partially occluding the blood flow in SVs. Chang *et al.* further proposed to incorporate the hemodynamic resistance of individual blood cells in the SVs toward the tail region (18). However, this zero-dimensional resistance modeling did not capture vascular remodeling-associated changes in flow rates and WSS. Given the complex geometrical features of the zebrafish micro-circulation, the variations in vessel diameter are pivotal for determining hemodynamics. For this reason, our CFD model addressed vasoconstriction and vasodilation, and vascular regeneration in the microvascular network, demonstrating a spatiotemporal-dependent modeling of vascular injury and repair *in vivo*.

Spatial and temporal variations in WSS modulate vascular endothelial function and homeostasis. WSS in the microcirculation regulate the canonical Wnt/β-catenin signaling pathway for vascular development (22, 36), and WSS-activated Notch signaling modulates endothelial loop formation and endocardial trabeculation (12, 20, 35). We have previously developed light-sheet imaging methods to study the development of trabeculation for zebrafish ventricles (10, 11). Combining light-sheet microscopy and computational fluid dynamics (CFD), we assessed the role of wall shear stress (WSS) in formation of trabeculation and outflow tract valvulogenesis in embryonic zebrafish (12–14). In this CFD modeling of vascular injury and repair, we further demonstrate the integration of CFD with genetic zebrafish system to uncover microvascular mechanics.

Blood effective viscosity provides the rheological basis to compute wall shear stress underlying cardiovascular morphogenesis. During development, blood viscosity regulates fluid shear stress to impart mechano-signal transduction to initiate cardiac trabeculation and outflow tract formation (12, 28). However, the minute amount of zebrafish blood renders measurements of blood viscosity experimentally challenging (37). We previously reported the capillary pressure-driven principle to develop microfluidic channels for measuring the small-scale blood viscosity (25), allowing for establishing a power-law non-Newtonian model to predict the transient rise and fall of viscosity during embryonic development. In this context, we further analyzed the non-Newtonian effects on the flow rates and wall shear stress in SVs by performing a parametric study based on the power-law index (*n*) (**Supplementary Fig. S1**). From the power law model, we normalized the results according to the reported value for non-Newtonian index (*n* = 1.09) (25). Our results indicate that despite local variations in WSS, the blood flow in micro=circulation was not significantly altered by variation of the non-Newtonian index. Overall, we unraveled the interplay between vascular remodeling and WSS in the face of tail amputation. Our CFD simulation was corroborated with micro PIV. The results of the study provide translational implications in micro-vascular injury and repair for human diseases with associated vascular complications, such as diabetes and peripheral artery disease.

## Acknowledgments

The present work was funded by the National Institutes of Health R01HL083015 (TKH), R01HL111437 (TKH), R01HL129727 (TKH), R01HL118650 (TKH), and VA MERIT AWARD I01 BX004356 (TKH).

## Notes

### Competing Interest Statement

The authors have declared no competing interest.

