## Supplementary Materials for "Computational Simulations of the 4-D Micro-Circulatory Network in Zebrafish Tail Amputation and Regeneration"

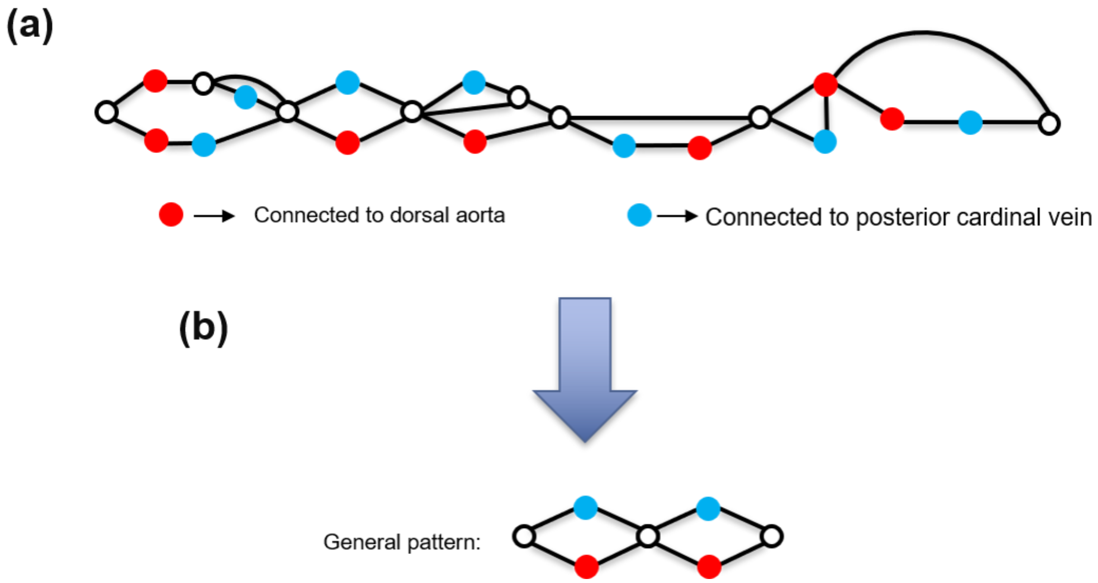


**Fig. S1.** (a) **The top-view from dorsal longitudinal anastomotic vessel (DLAV) network**. The pattern of connectivity between arterial and venous segmental vessels (SVs) was used to simulate the embryonic zebrafish micro-circulation. (b) **The propose generalized pattern of connectivity in DLAV network**. Note that the red circles denote the connection between arterial SV and DLAV, and the blues circles denote the connection between venous SV and DLAV.

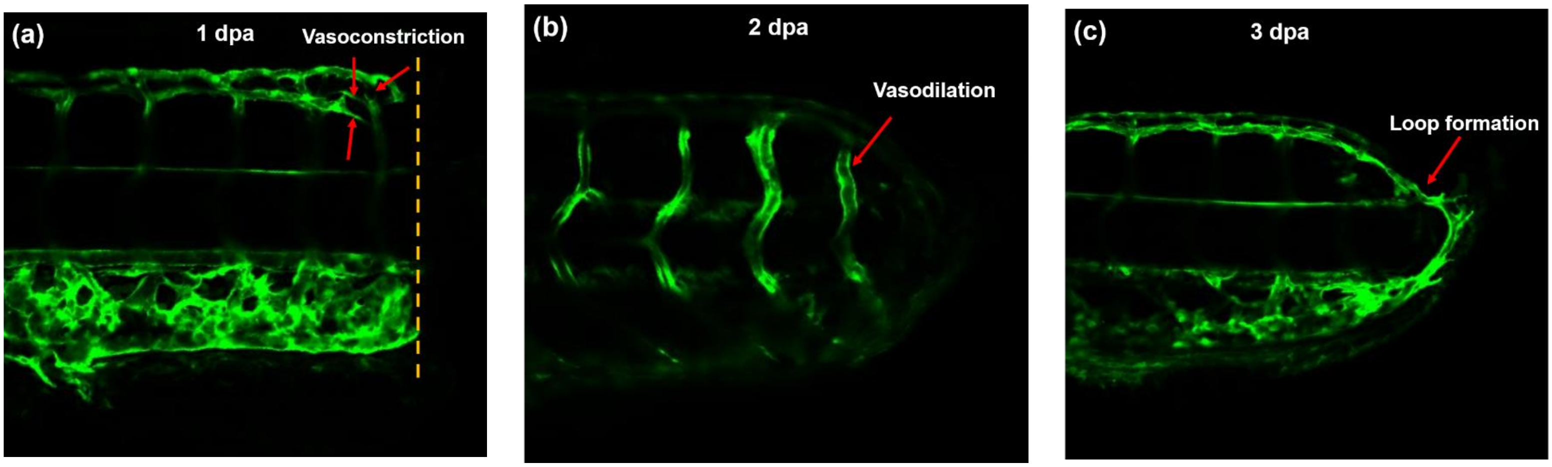


Fig. S2 Confocal images of green-fluorescent protein (GFP)-labeled endothelial cells post amputation. (a) Vasoconstriction proximal to amputation at DLAV-SV connection at 1 days post amputation (dpa). (b) Vasodilation in the SVs proximal to amputation at 2 dpa. (c) Regeneration of new vessel from the tips of arterial SV and PCV at 2 dpa.
